# Genomic structural variation in ‘Nebbiolo’ grapevines at the individual, clonal and cultivar levels

**DOI:** 10.1101/2020.10.27.357046

**Authors:** Simone Maestri, Giorgio Gambino, Andrea Minio, Irene Perrone, Emanuela Cosentino, Barbara Giovannone, Giulia Lopatriello, Luca Marcolungo, Dario Cantu, Marzia Rossato, Massimo Delledonne, Luciano Calderón

## Abstract

Structural Variants (SVs) are a widely unexplored source of genetic variation, both due to methodological limitations and because they are generally associated to deleterious effects. However, with the advent of long-range genomic platforms, it has become easier to directly detect SVs. In the same direction, clonally propagated crops provide a unique opportunity to study SVs, offering a suitable genomic environment for their accumulation in heterozygosis. In particular, it has been reported that SVs generate drastic levels of heterozygosity in grapevines. ‘Nebbiolo’ (*Vitis vinifera* L.) is a grapevine cultivar typical of north-western Italy, appreciated for its use in producing high-quality red wines. Here, we aimed to analyze the frequency of SVs in ‘Nebbiolo’, at three different organizational levels. For this purpose, we generated genomic data based on long-reads, linked-reads and optical mapping. We assembled a reference genome for this cultivar and compared two different clones, including *V. vinifera* reference genome (PN40024) in our comparisons. Our results indicate that SVs differentially occurring between ‘Nebbiolo’ clones might be rare, while SVs differentiating haplotypes of the same individual are as abundant as those that occur differentially between cultivars.

## Introduction

*Vitis vinifera* L. was one of the first crops to have a sequenced reference genome ^1^. In order to ease the assembly process due to the high levels of heterozygosity, inherent to grapevine cultivars, authors sequenced a nearly homozygous genotype derived from ‘Pinot Noir’ (PN40024) ^2^. Although technically convenient, focusing on homozygous materials has provided several limitations in understanding grapevines genomic complexity ^3^. As a clonally propagated crop, grapevines offer a suitable biological model for the study of structural variations (SVs), providing a proper genomic environment for SVs to accumulate as heterozygous recessives ^3^. In this regard, characterizing SVs in grapevines is a fundamental task, both because it has been shown they affect phenotypic aspects of domestication ^4^ and productive interest ^5^, and because they could be used as molecular markers for cost-effective clonal identification ^6^.

‘Nebbiolo’ (*V. vinifera* L.) is a prestigious grapevine cultivar used for high-quality red wines production (e.g. Barolo and Barbaresco); it has been cultivated since the 13th century in north-western Italy across the Piedmont, Aosta Valley and Lombardy regions ^6^. A recent study based on short-reads genomic data identified diagnostic single nucleotide variants (SNVs) among three clones, associated to different cultivation areas ^6^. It was also observed that ‘Nebbiolo’ estimated heterozygosity turned higher than the reported for other grapevine cultivars ^6^. This high heterozygosity and the availability of only short-reads data, contributed to make the *de novo* genome assembly and SVs identification in ‘Nebbiolo’ not possible at the time ^6^.

SVs are defined as genomic rearrangements of at least 50 bp in size ^7^, and include deletions (DELs), insertions (INSs) duplications (DUPs) and translocations (TRA) ^8^. Although the genome-wide characterization of SVs is considered essential for genome interpretation, progress on their study is notably lagging behind the thorough comprehension achieved for SNVs ^9,10^. In fact, widely adopted short-read platforms provide only indirect evidence to infer SVs presence ^11^, resulting in a high rate of SVs miscalls, especially in repetitive regions that short reads cannot resolve ^8^. However, a wide variety of long-range genomic platforms have recently emerged, including long-reads, optical mapping and linked-reads (i.e. synthetic long-reads), among many others ^9^. These new platforms have helped to overcome mapping issues in repetitive regions for example, thus enabling the direct detection of SVs ^9^.

At the same time, long-range genomic platforms have also contributed to improve the *de novo* genome assembly process, yielding highly contiguous assemblies up to chromosome-scale level. In particular, high-quality diploid genome assemblies for highly heterozygous crops have been obtained, such as: *Brassica rapa*, *Brassica oleracea*, *Manihot esculenta* ^4,12^, as well as several *V. vinifera* L. cultivars ^3,4,13–16^. These assemblies offer a smoother starting-point to identify novel features, previously hidden in collapsed and fragmented genome assemblies ^4,12^. Integration of multiple platforms is a common practice for obtaining high-quality genome assemblies ^2,3,12,14,17^. However, fewer studies have simultaneously compared the contribution of multiple platforms to detect SVs, especially in plants ^10,18,19^.

In this work we analyzed the occurrence of SVs across ‘Nebbiolo’ genomes by means of three different long-range genomic platforms, based on: long-reads, linked-reads, and optical mapping. We generated genomic data for two ‘Nebbiolo’ clones (CVT 71 and CVT 185), and in order to improve our ability to detect SVs we assembled a reference genome for this cultivar. Overall, we had the main purpose to compare the frequency of SVs at three different organizational levels: individual, clonal and cultivar. As a complementary objective, we evaluated the ability of the employed long-range genomic platforms at detecting SVs.

## Results

### *De novo* assembly of ‘Nebbiolo’ reference genome

We performed a *de novo* genome assembly for ‘Nebbiolo’ clone CVT 71, integrating PacBio (long-reads), Bionano Genomics (optical mapping) and Illumina (short-reads) data. First, we *de novo* assembled PacBio long-reads, yielding 875 primary contigs (767 Mbp) and 3,911 alternative haplotypes (i.e. haplotigs) (405 Mbp), with an assembly N50 = 1.2 Mbp. This preliminary assembly was polished with PacBio and Illumina reads. Then, we *de novo* assembled the Bionano single-molecule maps into 969 optical consensus maps totaling 1.1 Gbp (N50 = 1.5 Mbp); these maps were employed to anchor the polished preliminary assembly to produce the hybrid assembly. The hybrid assembly consisted of 978 anchored sequences (816 Mbp) and 4,000 not-anchored sequences (356 Mbp), totaling 1.2 Gbp (N50 = 2.5 Mbp), an assembly size which is twice as big as the expected haploid genome size. Therefore, we separated the two haplotypes to reduce redundancy, which may hamper SVs detection by decreasing mapping quality of reads aligned to homologous regions. This was performed by identifying shorter homologous sequences and moving them to a separate file, which we refer to as ‘Nebbiolo’ alternative haplotypes, as opposed to ‘Nebbiolo’ primary assembly. The obtained primary assembly was 561 Mbp in size, consisted of 230 sequences with a N50 = 5.4 Mbp, and included 94.8% of complete universal single-copy orthologue (BUSCO) genes. The obtained alternative haplotypes were 534 Mbp in size, consisted of 1,987 sequences with a N50 = 1.2 Mbp, and included 77.9% of complete BUSCO genes (**Table 1**). A total of 2,115 contigs totaling 107 Mbp (N50 = 0.06 Mbp) were discarded as they were identified as assembly artefacts.

**Table 1.** Contiguity and completeness statistics for ‘Nebbiolo’ CVT 71 genome assemblies. For BUSCO statistics, ‘C’ refers to gene completeness, ‘F’ to fragmented genes, and ‘M’ to missing genes.

|  | ‘Nebbiolo’<br>hybrid<br>assembly | ‘Nebbiolo’<br>primary<br>assembly | ‘Nebbiolo’<br>alternative<br>haplotypes |
| --- | --- | --- | --- |
| **Total assembly length (Mbp)** | 1,201.74 | 560.76 | 534.31 |
| **Assembly N50 (Mbp)** | 2.52 | 5.37 | 1.18 |
| **Total scaffolds length (Mbp)** | 716.22 | 487.55 | 228.67 |
| **Number of scaffolds** | 219 | 109 | 110 |
| **Number of gaps** | 655 | 395 | 260 |
| **Gaps size (Mbp)** | 29.22 | 20.52 | 8.70 |
| **Contigs in scaffolds** | 874 | 504 | 370 |
| **Remaining contigs** | 4,113 | 121 | 1,877 |
| **Remaining contigs total length (Mbp)** | 485.52 | 73.21 | 305.64 |
| **BUSCO statistics** | C:96.7%,<br>F:1.1%,<br>M:2.2% | C:94.8%,<br>F:2.0%,<br>M:3.2% | C:77.9%,<br>F:1.9%,<br>M:20.2% |

### Heterozygous structural variations are abundant at the individual level

Genomic data for ‘Nebbiolo’ CVT 71 was aligned to the ‘Nebbiolo’ primary assembly to identify heterozygous polymorphisms across CVT 71 genome (**Table 2**).

**Table 2.** Raw genomic data obtained for ‘Nebbiolo’ clones CVT 71 and CVT 185.

| ‘Nebbiolo’<br>clone | Sequencing<br>platform | Data type | N50 reads/<br>molecules<br>(bp) | Number of<br>reads/molecules<br>generated | Number of<br>bases<br>generated<br>(Gbp) |
| --- | --- | --- | --- | --- | --- |
| CVT 71 | PacBio | Long-reads | 27,197 | 3,286,690 | 51 |
| CVT 71 | Bionano<br>Genomics | Single-molecule<br>maps | 241,361 | 654,883 | 157 |
| CVT 71 | 10x Genomics | Linked-reads | 2x150 | 273,179,528 | 82 |
| CVT 185 | 10x Genomics | Linked-reads | 2x150 | 261,527,174 | 78 |

Mean coverage reached with each set of genomic data was 64X (PacBio), 63X (10x Genomics linked-reads) and 77X (Bionano Genomics). The sensitivity of each platform at detecting SVs, after filtering, was quite different. PacBio identified the highest number of SVs (6,760), followed by 10x Genomics (2,625), and Bionano Genomics (657) (**Figure 1** and **Table 3**).

**Figure 1.**
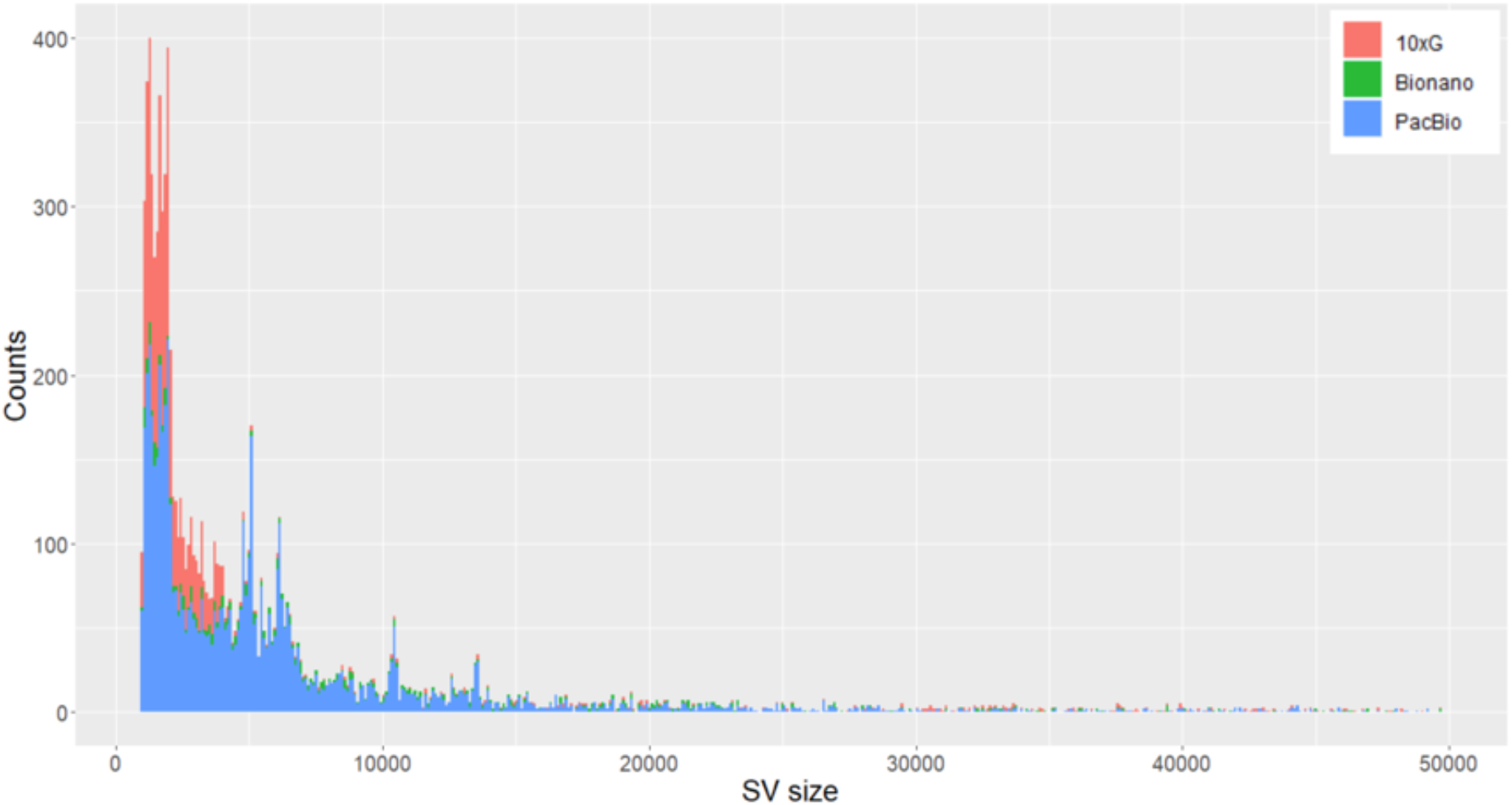
Size distribution of Structural Variants (SVs) within ‘Nebbiolo’ CVT 71 identified by PacBio, 10x Genomics and Bionano Genomics. For each SV size, a colored bar represents the number of SVs identified by each platform.

**Table 3.** Structural Variant (SV) types identified comparing ‘Nebbiolo’ CVT 71 haplotypes.

| Sequencing platform | DELs | INSs/DUPs | INVs | TOT | Cumulative size of SVs (Mbp) |
| --- | --- | --- | --- | --- | --- |
| PacBio | 5,951 | 722 | 87 | 6,760 | 101 |
| 10x Genomics | 2,546 | 19 | 12 | 2,625 | 55 |
| Bionano genomics | 331 | 326 | 0 | 657 | 31 |

Cumulative size of SVs identified by PacBio data represents 18.1% of the primary genome assembly, indicating that heterozygous SVs are abundant at the individual level. In particular, deletions/insertions appeared as the most abundant type of SV detected, suggesting that a great proportion of ‘Nebbiolo’ genome is unbalanced (i.e. hemizygous) (**Table 3**). Out of all SVs identified by PacBio, 46% overlapped to SVs identified by 10x Genomics, and 5% overlapped to SVs identified by Bionano Genomics maps. While, out of all SVs identified by 10x Genomics, 82% overlapped to SVs identified by PacBio; and out of all SVs identified by Bionano Genomics only 31% overlapped to SVs identified by PacBio (**Figure 2**).

**Figure 2.**
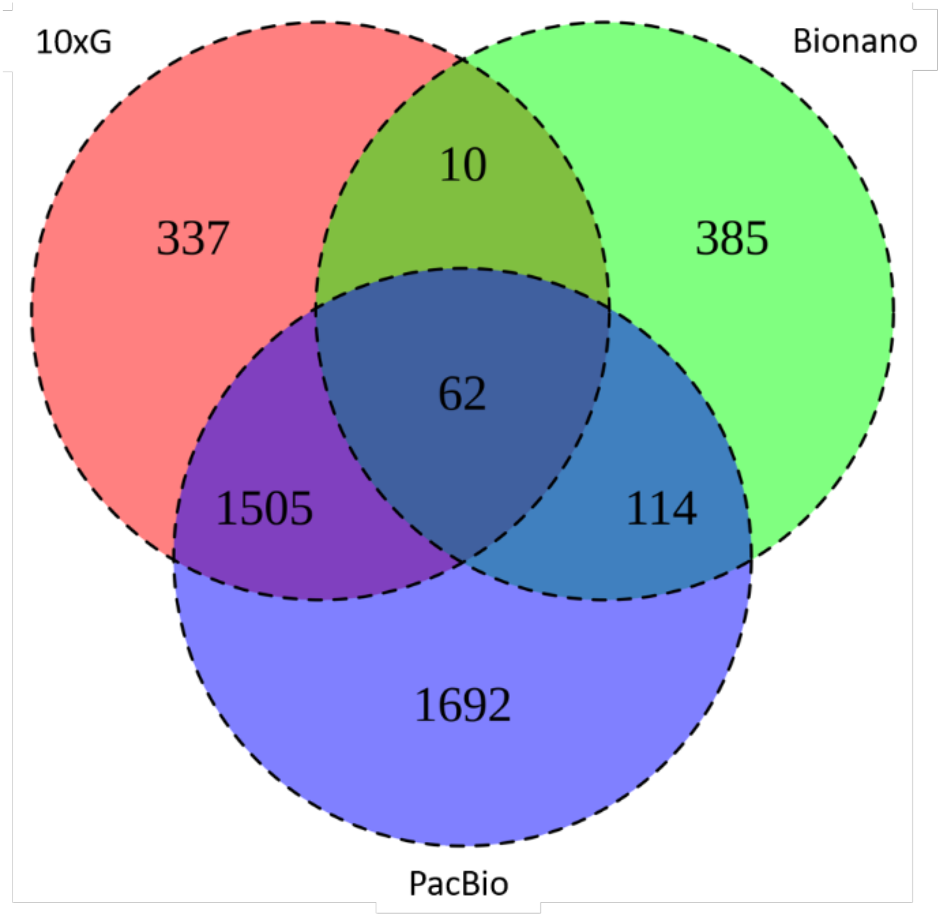
Intersection of Structural Variants (SVs) within ‘Nebbiolo’ CVT 71 identified by PacBio, 10x Genomics and Bionano Genomics. SVs identified by each platform are intersected based on the genomic coordinates at which they are called.

Manual inspection of a subset of PacBio and 10x Genomics reads alignments confirmed all SVs called using PacBio data as true variants. Moreover, many SVs called only by PacBio were supported also by 10x Genomics reads, despite not having been identified by the variant caller (e.g. **Figure S1**).

### No evidence of structural variations between ‘Nebbiolo’ clones CVT 71 and CVT 185

10x Genomics linked-reads obtained for both clones were aligned to the ‘Nebbiolo’ CVT 71 primary assembly to identify structural polymorphisms between clones. Mean coverage values were 63X (CVT 71) and 60X (CVT 185). After performing SVs calling and quality filtering, a total of 2,651 SVs were identified for clone CVT 185 and 2,625 SVs for CVT 71 (**Table 4**).

**Table 4.** Structural Variant (SV) types identified by 10x Genomics comparing ‘Nebbiolo’ CVT 71 and CVT 185 clones.

| ‘Nebbiolo’ clone | DELs | INSS/DUPs | INVs | TOT | Cumulative size of SVs (Mbp) |
| --- | --- | --- | --- | --- | --- |
| CVT 71 | 2,546 | 19 | 12 | 2,625 | 55 |
| CVT 185 | 2,584 | 11 | 6 | 2,651 | 49 |

The intersection of the two sets of variants suggested that 321 SVs occurred only in clone CVT 185 and not in CVT 71 (**Figure 3**).

**Figure 3.**
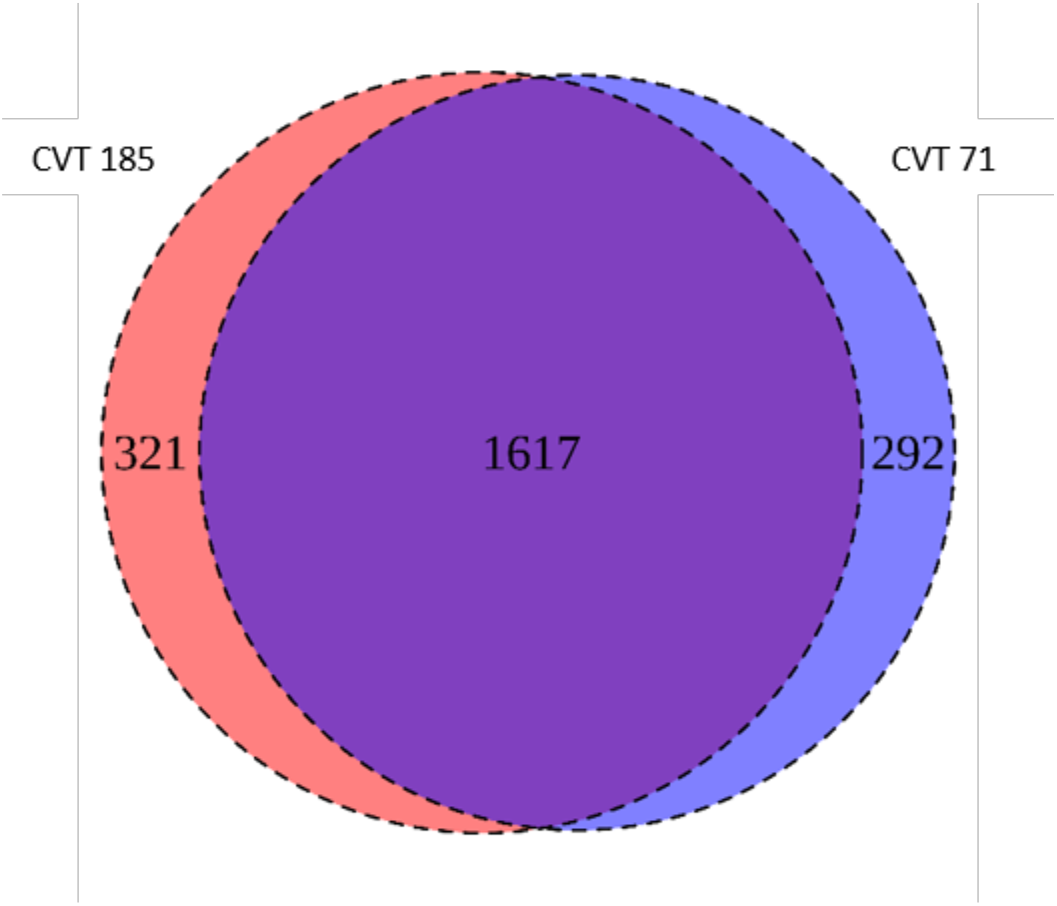
Intersection of Structural Variants (SVs) identified by 10x Genomics occurring between ‘Nebbiolo’ clones CVT 71 and CVT 185. The bioinformatic analysis shows the presence of 321 SVs occurring in CVT 185 clone that are not shared with clone CVT 71.

However, none of the latter variants were confirmed as occurring in only one clone after manual inspection. We observed that, even when an SV was called exclusively for one clone, the manual inspection revealed similar number of reads supporting that SV for both clones (e.g. **Figure S2**).

### Structural variations between cultivars are as frequent as at individual level

We investigated SVs at the cultivar level comparing ‘Nebbiolo’ genomic data to *V. vinifera* reference genome PN40024 12X v2 ^20^. After alignment, mean coverage values were 70X (PacBio), 72X (10x Genomics) and 38X (Bionano Genomics). After performing variant calling analyses and filtering, each platform was able to identify different numbers of SVs: 7,808 (PacBio), 2,381 (10x Genomics), and 1,039 (Bionano) (**Figure 4** and **Table 5**).

**Figure 4.**
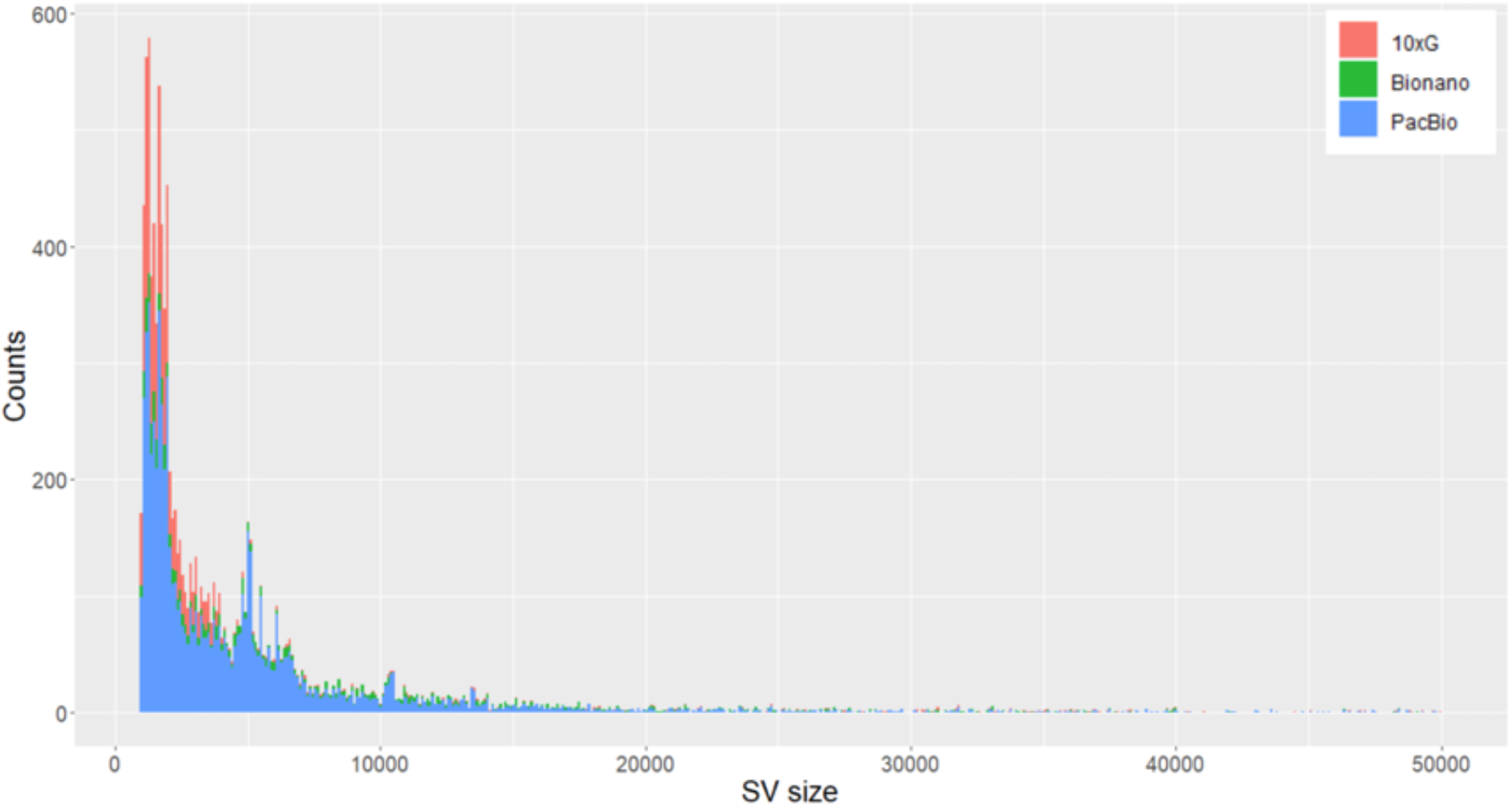
Size distribution of Structural Variants (SVs) between ‘Nebbiolo’ CVT 71 and PN40024 identified by PacBio, 10x Genomics and Bionano Genomics. For each SV size, a colored bar represents the number of SVs identified by each platform.

**Table 5.** Structural Variant (SV) types identified comparing ‘Nebbiolo’ CVT 71 to PN40024.

| Sequencing platform | DELs | INs/DUPs | INVs | TOT | Cumulative size of SVs (Mbp) |
| --- | --- | --- | --- | --- | --- |
| PacBio | 6,565 | 1,115 | 128 | 7,808 | 47 |
| 10x Genomics | 2,352 | 0 | 7 | 2,381 | 9 |
| Bionano genomics | 279 | 757 | 3 | 1,039 | 21 |

The number of SVs identified by PacBio and 10x Genomics did not change drastically, compared to the number of SVs identified at individual level (**Tables 3** and **5**). However, when making the same comparison, SVs detected by Bionano Genomics almost doubled (**Table 5**), possibly due to better anchoring of consensus maps to a chromosome-scale assembly. Nonetheless, the cumulative size of SVs almost halved with respect to SVs identified at individual level, most likely due to several large SVs that were discarded, as consequence of the presence of 15,325 gaps (15.99 Mbp) across PN40024 genome (**Table S1**).

Out of all SVs identified by PacBio, 42% overlapped to SVs identified by 10x Genomics, and 10% overlapped to SVs identified by Bionano Genomics. While, out of all SVs identified by 10x Genomics, 88% overlapped to SVs identified by PacBio; and out of all SVs identified by Bionano Genomics, 43% overlapped to SVs identified by PacBio (**Figure 5**).

**Figure 5.**
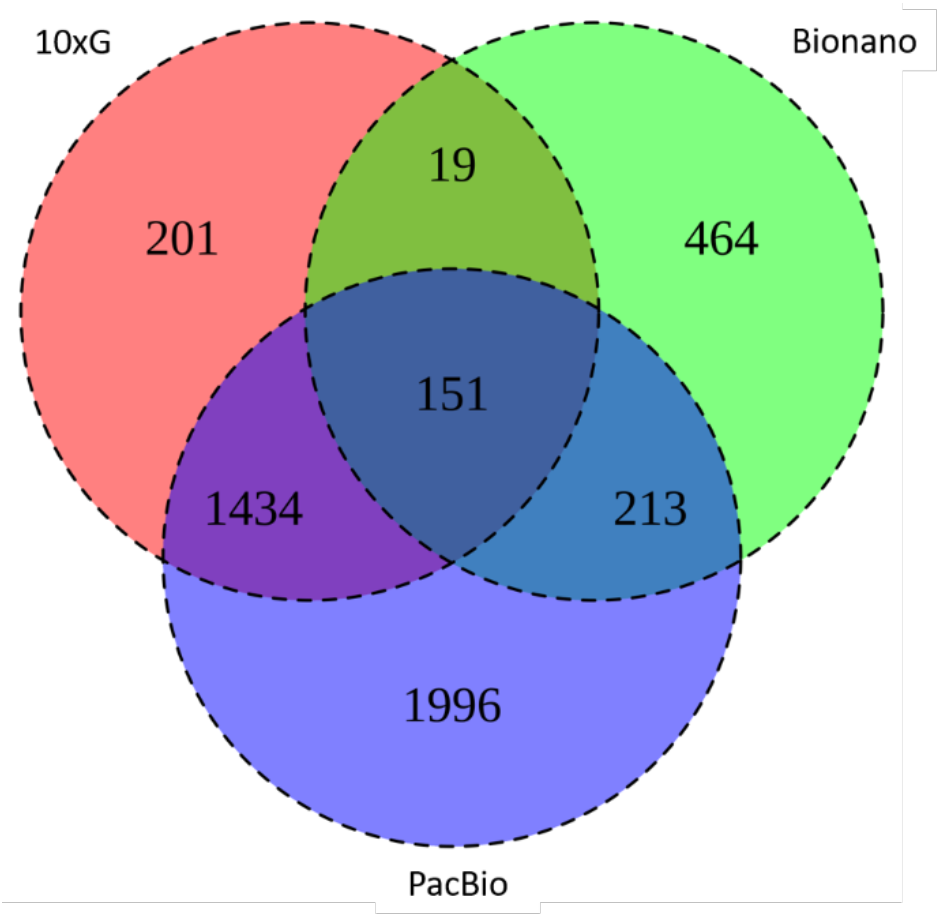
Intersection of Structural Variants (SVs) between ‘Nebbiolo’ CVT 71 and PN40024 identified by PacBio, 10x Genomics and Bionano Genomics. SVs identified by each platform are intersected based on the genomic coordinates at which they are called.

Manual inspection confirmed our previous observations, any given SV called by PacBio is reliable regardless of the other platforms supporting that same SV (e.g. **Figure S3**).

## Discussion

Significant innovations in genomic platforms have recently enabled a deeper comprehension of plant genomes complexity, allowing the obtainment of phased genome assemblies and SVs identification ^2,12^. In particular, a detailed characterization of both haploid complements of the diploid genome of some grapevine cultivars has been recently achieved ^3,13,15^.

We aimed to detect SVs across Nebbiolo’s highly heterozygous genome by generating genomic data with three long-range platforms. In order to maximize our ability to detect SVs, we assembled *de novo* a reference genome for ‘Nebbiolo’ (clone CVT 71). The size of the obtained primary assembly (561 Mbp) turned 15% bigger than the partially inbred ‘Pinot Noir’ PN40024 (486 Mbp) ^20^ and ‘Chardonnay’ (490 Mbp) ^21^ assemblies. At the same time, our assembly resulted smaller than those of other *V. vinifera* cultivars, e.g. ‘Cabernet Sauvignon’ (590 Mbp) ^15^, ‘Zinfandel’ (591 Mbp) ^16^, ‘Chardonnay’ (605 Mbp) ^3^ and ‘Carménère’ (623 Mbp)^13^. Variations in the primary assembly size might be mainly explained in terms of retention of both copies of some heterozygous regions in the primary assembly ^13^. In fact, despite all the mentioned assemblies were obtained using diploid-aware FALCON-Unzip pipeline, only the one described in Roach et al. ^21^ was processed with bioinformatic tools aimed at optimizing reassignment of allelic contigs, based on homology and read-coverage ^22^. Supporting this observation, two assemblies for ‘Chardonnay’ cultivar, that differ in the latter methodological aspect, were reported ^3,21^, showing 20% difference in size. Here, primary contigs produced by Falcon-Unzip resulted 37% bigger than the final primary assembly. This means that high heterozygosity makes it challenging to correctly discriminate between primary and alternative contigs, leading to imprecise estimates of the haploid genome size. In our case, long-range information provided by optical consensus maps proved to be valuable for scaffolding contigs that come from the same haplotype, increasing the N50 value of 67%. This allowed to improve the contiguity and to reduce haplotype switch errors, smoothing the identification of homologous regions.

We took advantage of comparisons at the individual and cultivar levels to test the ability of each employed long-range genomic platform at detecting SVs. Long-reads turned to be the platform identifying the highest number of reliable SVs, followed by linked-reads and by optical mapping. Considering SVs called by long-reads as the ‘gold standard’, linked-reads proved to be almost as precise although not as sensitive. On the other hand, optical mapping data turned to be the least precise and sensitive for SVs detection. These results suggest that linked-reads may represent a cost-effective alternative to long-reads, for example in population-based studies ^23^. We also observed that, like with short-reads, deletions (DELs) were the most abundant type of SVs detected, while insertions (INSs) were detected in lower number. This could be explained by the higher algorithmic difficulty of calling insertions through mapping approaches ^9^, and from the stringent filtering criteria we applied by keeping only SVs for which precise breakpoints coordinates could be determined. The high deletions/insertions ratio suggest that the estimates about the extent of heterozygous SVs in ‘Nebbiolo’ are conservative, as the same SV could be seen as a DEL or an INS depending on which haplotype is included in the primary assembly. Without any algorithmic bias, the number of heterozygous INSs should approach that of heterozygous DELs.

To explore the frequency of SVs in ‘Nebbiolo’ grapevines at different organizational levels we performed the analysis at individual level, between two clones and between two cultivars. Our analyses at the individual level showed that a significant proportion of ‘Nebbiolo’ genome is affected by SVs that differentiate the two haplotypes. In particular, the cumulative size of SVs identified by long-reads represents 18.1% of the primary assembly size, a value considerably higher than the 6.94% reported for ‘Zinfandel’ 16, but similar to the 15.1% reported for ‘Chardonnay’ 3. This finding supports the observation that hemizygosity is rampant in clonally propagated grapevine genomes ^3^, and also the previously proposed high diversity between ‘Nebbiolo’ parental haplotypes ^6^. Few studies have performed a genome-wide analysis of somatic mutations accumulated in different clones of the same cultivar ^6,16,21^. Among these, only Vondras et al. ^16^ studied the distribution of SVs among ^15^ ‘Zinfandel’ clones, based solely on short-reads genomic data, and they observed that SVs are rarer than SNVs. Even though thousands of SVs were reported among ‘Zinfandel’ clones, authors pointed out that additional work should be undertaken to confirm these variants ^16^. Our results add further evidence to the following concepts: firstly, a thorough validation is essential for spotting spurious SV calls ^5,18,23^. In our case, none of the 321 SVs that were supposed to occur in one clone and not in the other, passed the manual inspection criteria. Secondly, as observed for ‘Zinfandel’, SVs differentiating ‘Nebbiolo’ clones are clearly less frequent than SNVs ^6^.

In regard to the comparisons at the cultivar level (‘Nebbiolo’ vs PN40024), we observed that the total number of SVs identified was similar to that observed at the individual level. A reason for the latter could be that all extant cultivars originated from out-crossing two pre-existing cultivars and have been preserved through clonal propagation ^24^. The abundancy of heterozygous SVs likely originated from the pre-existing genetic differences between parental cultivars; while clonality makes possible to maintain, even to increase, this heterozygosity. On the other hand, the cumulative size of SVs at the cultivar level almost halved the observed at individual level, probably as a by-product of assembly issues related to gaps abundancy across PN40024 reference genome. Large SVs are more likely to overlap a gap, this provokes their removal during the filtering process causing the observed difference in the cumulative size. This provides further evidence that using a high-quality genome assembly is a fundamental prerequisite to fully understand the genomic complexity of the focal organism. In particular, increasing the contiguity of an assembly will turn into a more precise SVs calling process.

In conclusion, we found that SVs differentiating clones of the same cultivar might be rare in grapevines, and that SVs seem to occur at the individual level with a similar frequency than between cultivars.

## Methods

### Samples collection and genomic data generation

Samples to assemble the *V. vinifera* L. cv. ‘Nebbiolo’ reference genome correspond to clone CVT 71. The material was sampled at IPSP CNR, Turin. For PacBio sequencing, DNA was extracted at the Functional Genomics Laboratory (University of Verona, Italy) from 1 g of young leaves of ‘Nebbiolo’ CVT 71 using the cetyltrimethylammonium ammonium bromide (CTAB) extraction buffer ^25^ modified from Japelaghi et al. ^26^ and Healy et al. ^27^, as described in Chin et al. ^15^ and combined with PacBio Guidelines for gDNA clean up. The purity of extracted DNA was assessed using NanoDrop™ 1000 Spectrophotometer (Thermo Scientific, Germany). Genomic DNA concentration was fluorometrically measured combining dsDNA Broad Range Assay Kit with Qubit^®^ 4.0 (Thermo Fisher Scientific, Waltham, USA); the size of DNA fragments was evaluated using the CHEF Mapper electrophoresis system (Bio-Rad Laboratories, California). Genomic DNA (16 μg) was used to prepare a single-molecule real-time (SMRT) bell library according to the manufacturer’s protocol (Pacific Biosciences; 30-kb template preparation using BluePippin (SageScience) size selection system with a 20-kb cut-off). Sequencing was performed on a PacBio RS II platform (Pacific Biosciences, CA, USA) producing 3,286,690 reads with a N50 of 27,179 bp and a total of 51 Gbp of SMRT data using PacBio P6-C4 chemistry. Library preparation and sequencing were performed at the University of California Davis (California, USA).

Bionano Genomics mapping is based on the enzymatic digestion of high-molecular weight DNA molecules by a nicking enzyme, followed by the incorporation in the nicks of a fluorescent nucleotide. Labelled molecules are scanned and distances between labels are recorded after image digitalization ^28^. Here, we used young grapevine plants of ‘Nebbiolo’ clone CVT 71, expanded *in vitro* on solid sterile culture media. High Molecular Weight DNA was extracted from 1 g of freshly harvested leaves using the IrysPrep Plant Tissue DNA Isolation Protocol (Bionano) with minor adjustments as described in Cecchin et al. ^17^. The Mega-base size of extracted DNA was verified by Pulsed-Field-Electrophoresis (PFGE). DNA (510 ng) was labelled and stained using 3.4 μl of Nb.BssSI (20U/μl) nicking endonuclease in combination with the NLRS DNA labelling kit (Bionano Genomics) and the labelled DNA was loaded on an Irys chip, generating 157 Gbp of molecules. DNA extraction, labeling and image acquisition were performed at the Functional Genomics Laboratory (University of Verona, Italy).

For 10x Genomics library preparation of ‘Nebbiolo’ clones CVT 185 and CVT 71, high-molecular-weight DNA was extracted from a nuclear preparation obtained from 1 g of young leaves. Tissue grounded in liquid nitrogen was resuspended in NIBTM (10 mM Tris pH 8, 10 mM EDTA pH 8, 0.5 M Sucrose, 80 mM KCl, 8% (w/v) PVP-10, 100 mM Spermine, 100 mM Spermidine, pH 9.0) supplemented with 0.5% Triton-100 and 0.2% beta-mercaptohetanol and kept on ice for 30 min. The tissue homogenate was filtered first through a 100-μm and then through a 40-μm cell strainer, then centrifuged at 2500 g for 20 min at 4 °C in a swing bucket rotor. Nuclei pellet was resuspended gently and washed with 30 ml of cold buffer and spun at 60 g for 2 minutes at 4 °C with no deceleration to remove tissues debris. The supernatant containing nuclei was filtered through at 40-μm cell strainer and spun to pellet the nuclei again at 2500 g for 20 min. The latter step was repeated until a white nuclei pellet and a clear supernatant were obtained. DNA was extracted from the isolated nuclei pellets using the Qiagen Genomic tip-100 (Qiagen) following the manufacturer’s instructions. Size of extracted DNA was verified by Pulsed-Field-Electrophoresis (PFGE). A 10x GEM library was constructed according to manufacturer’s recommendations (10x Genomics) starting from 10ng HMW DNA. DNA extractions and library preparations were performed at the Functional Genomics Laboratory (University of Verona, Italy). Libraries were quantified by qPCR and sequenced at Macrogen Inc. (Seoul, South Korea), using an Illumina HiSeq X Ten instrument. Sequencing produced 273,179,528 and 261,527,174 paired-end reads with length 150 bp and median insert size 315 bp and 326 bp for CVT 71 and CVT 185 clones respectively (**Table 2**). Mean molecule length of 10x Genomics barcoded libraries were 85,986 bp and 69,603 bp for CVT 71 and CVT 185 clones respectively.

### *De novo* Genome assembly and scaffolding of the ‘Nebbiolo’ genome

A preliminary *de novo* assembly based on PacBio long-reads from CVT 71 clone was performed using FALCONUnzip-DClab ^13^. This is a pipeline based on FALCON Unzip v1.7.7 15 and Damasker v1.0p1 ^29^, available at https://github.com/andreaminio/FalconUnzip-DClab. In detail, repeats were first masked using TANmask and REPmask modules in Damasker; reads were then corrected with Falcon Correct and repeats were masked also on corrected reads; afterwards, reads were assembled using FALCON. Multiple parameters were tested to produce the least fragmented assembly; haplotype reconstruction of the best assembly was then performed with Falcon-Unzip, producing a set of 875 primary contigs (767 Mbp) and 3,911 alternative haplotypes (405 Mbp). The preliminary assembly was 1.2 Gbp in size, with a N50 = 1.2 Mbp. Polishing of the preliminary assembly was performed using Arrow algorithm from GenomicConsensus v2.3.3. package (Pacific Biosciences). Illumina data for clone CVT 71 produced elsewhere 6 was used to polish the assembly. Illumina reads were mapped to the preliminary assembly with BWA mem v0.7.17 ^30^ and the resulting ·bam file was used for polishing with Pilon v1.23 ^31^. Bionano optical maps were employed to correct mis-assemblies by anchoring NGS contigs to consensus maps with Bionano Solve v3.4 hybrid scaffolding pipeline, with the following settings: *expected genome size of 0.5 Gbp, no preassembly, cut CMPR, non-haplotype, extend and split, cut segdups and Irys instrument*. In particular, single-molecule maps were assembled into 969 consensus maps totaling 1.1 Gbp, with a N50 = 1.5 Mbp. Consensus maps were used to produce a diploid hybrid assembly composed of 978 anchored sequences (816 Mbp) and 4,000 not-anchored sequences (356 Mbp). The hybrid assembly was 1.2 Gbp in size, with a N50 = 2.5 Mbp. To reduce redundancy and separate the primary assembly from the alternative haplotypes, two rounds of purging were performed with purge_haplotigs v1.1.0 22, yielding a primary assembly composed of 230 sequences (561 Mbp) with a N50 = 5.4 Mbp and 1,987 alternative haplotypes (534 Mbp) with a N50 =1.2 Mbp. BUSCO v4.1.2 ^32^ was applied to the assembly using the *eudicotyledons_odb10* database, to assess genome completeness. The obtained primary assembly (561 Mbp) was used as the ‘Nebbiolo’ reference for SVs calling. Assembly statistics were calculated with assembly-stats v1.0.1 (https://github.com/sanger-pathogens/assembly-stats).

### Structural Variants identification

SV calling was performed after aligning either reads or consensus genome maps to the corresponding reference genome. More precisely, for comparisons at the individual (i.e. between haplotypes) and clonal levels (CVT 71 vs. CVT 185), the obtained ‘Nebbiolo’ primary assembly was employed as the reference genome. For comparisons at the cultivar level, *V. vinifera* PN40024 12X v2 primary assembly was employed as reference genome. PacBio long reads were aligned to the reference using NGMLR v0.2.7 ^23^. SVs were called using Sniffles v0.1.11 ^23^ and we removed all SVs with the IMPRECISE and non-PASS flags. Single-molecule maps produced by Bionano Genomics for clone CVT 71 were aligned to the reference and assembled de novo into consensus genome maps, using Bionano Solve v. 3.4 with the following settings: *haplotype, expected genome size of 0.5 Gbp, no preassembly, cut CMPR* and *Irys instrument*, obtaining a consensus genome maps N50 = 1.3 Mbp. These maps were aligned to the ‘Nebbiolo’ reference genome to identify inconsistencies representing SV events occurring between consensus genome maps and the reference. Bionano Genomics key file was used to revert anchors names to original chromosomes names, to enable further comparisons. 10x Genomics reads were aligned to the reference and SVs were called using Longranger v2.2.2, variants with non-PASS flag were discarded. SVs identified by the three platforms overlapping to regions containing ambiguous nucleotides were removed with Bedtools intersect v2.28.0 ^33^. This was performed after the observation that many spurious SVs were called overlapping to gaps or hard-masked regions. Retained SV calls were merged using SURVIVOR merge v1.0.7 ^34^, setting 50 kbp as the maximum allowed distance between breakpoints. This permissive threshold was chosen, in order not to consider two SVs as different events due to imprecise breakpoint detection of Bionano Genomics platform ^35^. Translocations were excluded from all analyses because they turned technically difficult to process through our bioinformatic pipelines. Since each variant caller represented them in a different way, either as a pair of breakpoints with SVTYPE=BND (Sniffles and Longranger) or as a single event with SVTYPE=TRA (Bionano Solve). We also removed all SVs < 1 kbp from the variants list, because Bionano Genomics platform is not able to accurately detect SVs shorter than 500 bp ^35^. The sizes of the final set of SVs identified by the three platforms were plotted in R v3.6.0 with ggplot2 package ^36^. The cumulative size of SVs was calculated adding up the absolute sizes of the retained SVs. Manual inspection was performed with IGV genome browser v2.4.17, and the criteria to select the inspected SVs was based on the obtained Venn diagrams. More precisely, we randomly chose for manual inspection three SVs for each of the subsets identified by each platform, both at the individual (**Figure 2**) and cultivar levels (**Figure 5**), totaling 42 SVs. At the same time, with the aim to identify clone specific variants, all 321 SVs called only for clone CVT 185 (**Figure 3**) were manually inspected. SVs were considered as validated if they were supported by at least four non-reference reads.

## Data availability

The V. vinifera ‘Nebbiolo’ genome project data were deposited at NCBI under BioProject XXX and BioSample YYY. Raw 10x Genomics sequenced reads for clones CVT 71 and CVT 185 were deposited in the NCBI Sequence Read Archive under accession number ZZZ.

## Acknowledgments

We are grateful for the technical assistance of Rosa Figueroa-Balderas. DC was partially supported by NSF grant no. 1741627, E. & J. Gallo Winery, and the Louis P. Martini Endowment in Viticulture. LC staying at University of Verona to conduct this project was possible thanks to grants awarded by IILA-Organizzazione internazionale italo-latino americana and University of Verona.

## Authors Contributions

S.M. performed most of the bioinformatic and statistical analyses and wrote the manuscript. D.C. and A.M. generated the PacBio genomic data and the corresponding *de novo* genome assembly. G.G., I.P., E.C, B.G and, M.R. generated Illumina, 10X and Bionano data. L.M. and G.L. collaborated with the bioinformatic analysis. M.D. designed the project and structured the manuscript. L.C. coordinated the project, collaborated with the bioinformatic analyses and wrote the manuscript. All authors read and provided valuable advice to improve this manuscript.

