## Supplementary figures for "Genomic structural variation in ‘Nebbiolo’ grapevines at the individual, clonal and cultivar levels"

**Figure S1. IGV screenshot of a validated heterozygous SV in Nebbiolo CVT 71 genome.**

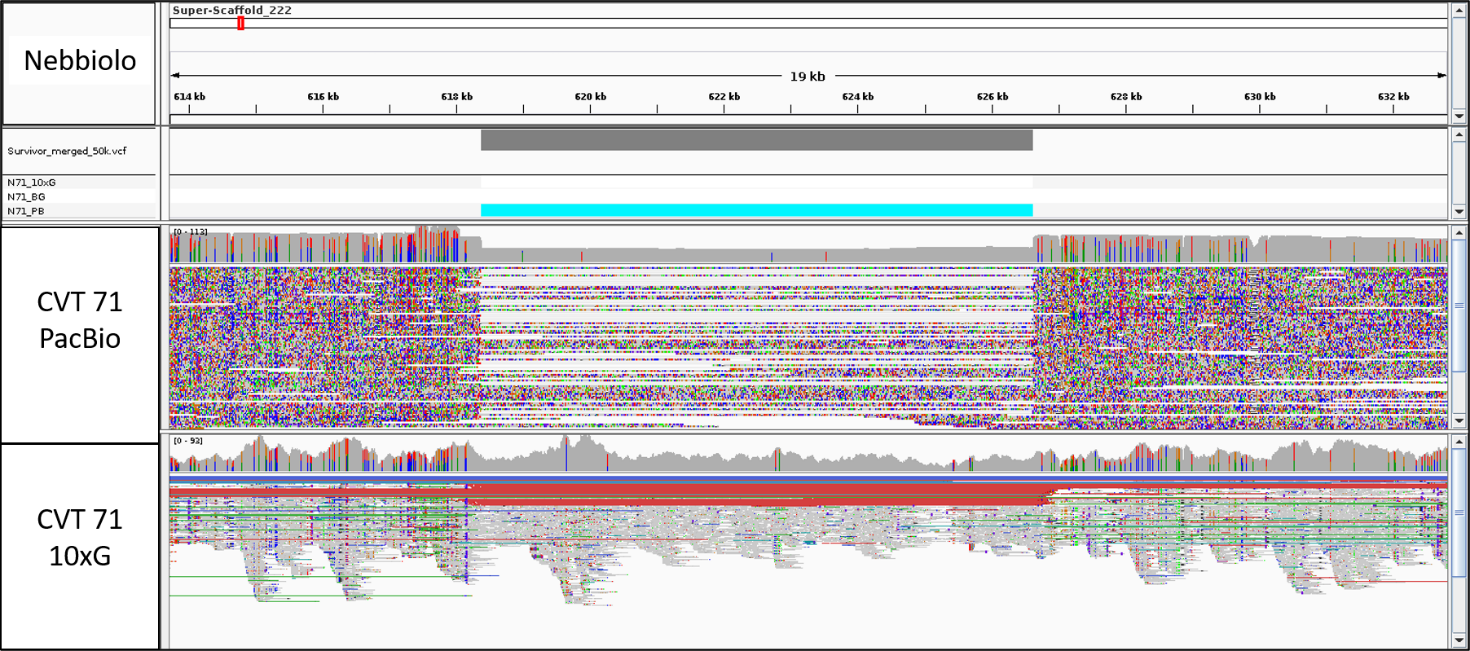

**Figure S2. IGV screenshot of a not validated SV between Nebbiolo clones CVT 71 and CVT 185.**

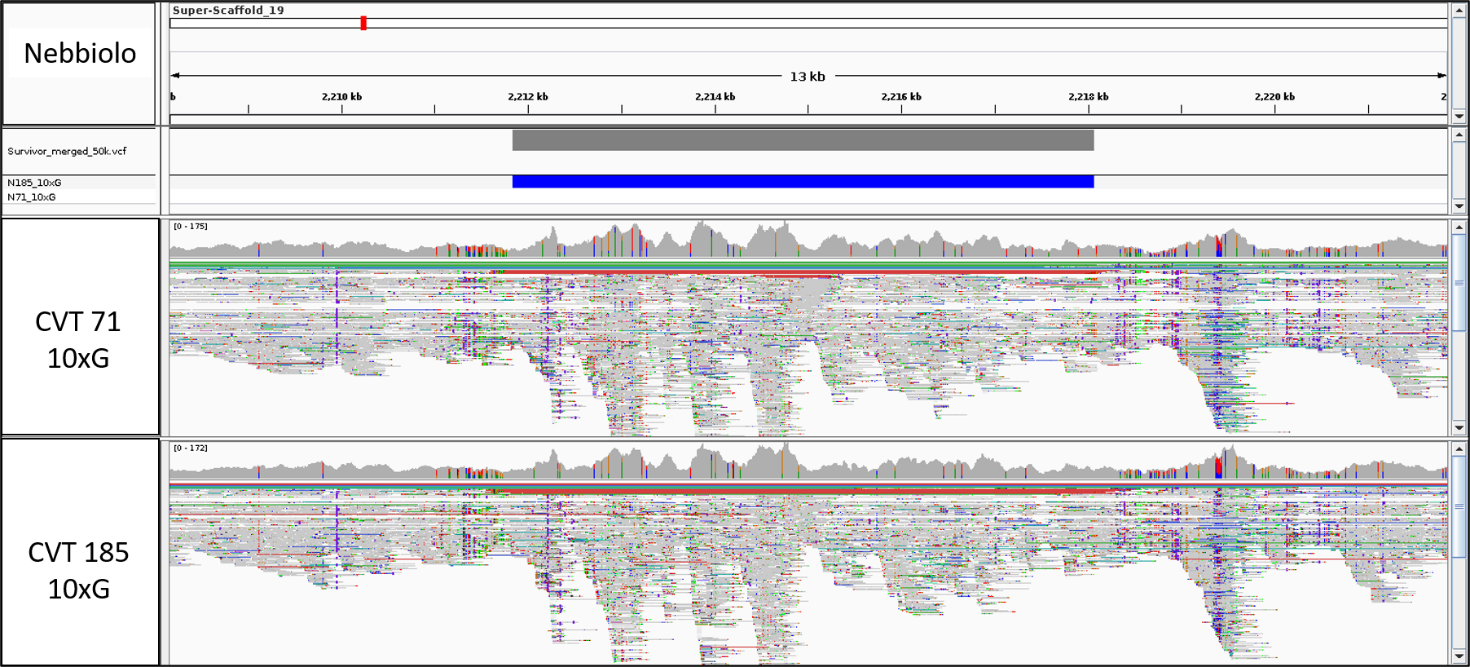

**Figure S3. IGV screenshot of a validated SV between Nebbiolo CVT 71 and PN40024.**

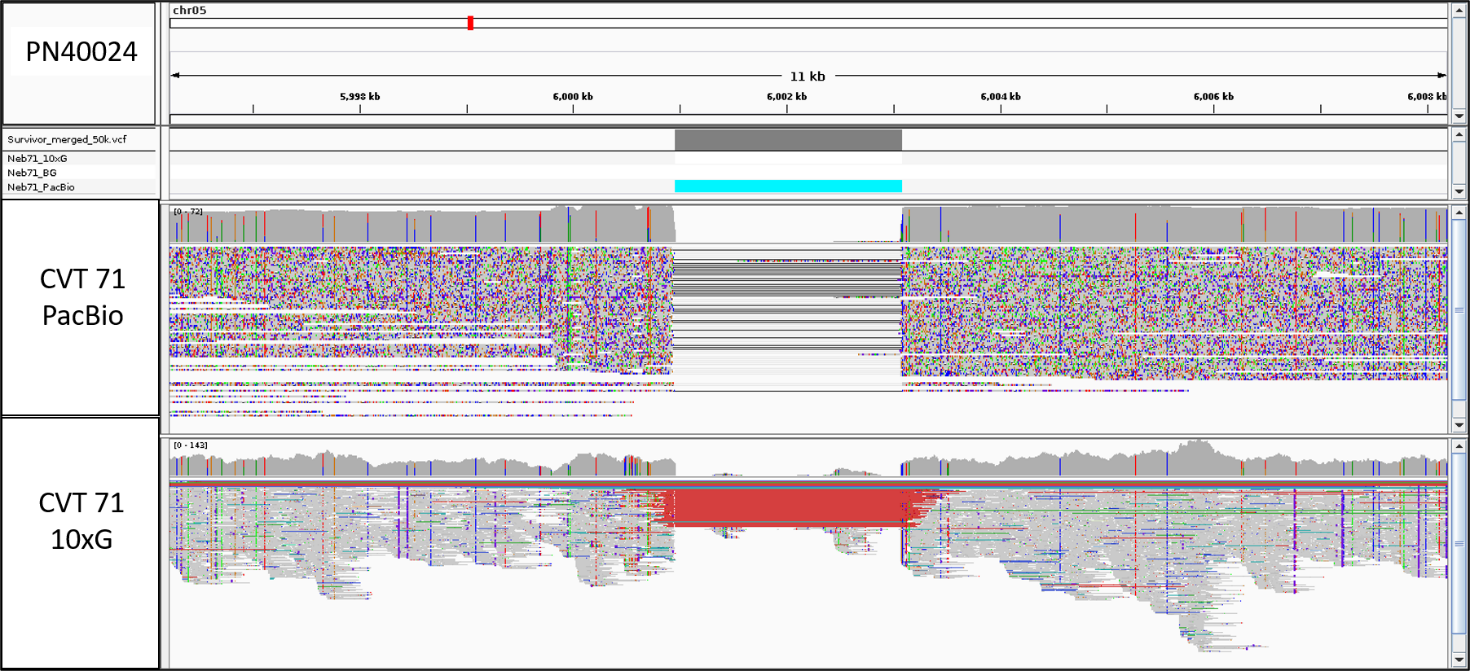

**Supplementary Tables**

**Table S1. Contiguity and completeness statistics for PN40024 V2 genome assembly.** For BUSCO statistics, ‘C’ refers to gene completeness, ‘F’ to fragmented genes, and ‘M’ to missing genes.

|  | **PN40024 V2 assembly** |
| --- | --- |
| **Total assembly length (Mbp)** | 486.21 |
| **Assembly N50 (Mbp)** | 24.27 |
| **Number of scaffolds** | 20 |
| **Number of gaps** | 15,325 |
| **Gaps size (Mbp)** | 15.99 |
| **BUSCO statistics** | C:96.0%, F:1.4%, M:2.6% |
